# Integration of Cell Growth and Asymmetric Division During Lateral Root Initiation In *Arabidopsis thaliana*

**DOI:** 10.1101/2020.12.15.422941

**Authors:** Lilli Marie Schütz, Marion Louveaux, Amaya Vilches Barro, Sami Bouziri, Lorenzo Cerrone, Adrian Wolny, Anna Kreshuk, Fred A. Hamprecht, Alexis Maizel

**Affiliations:** Center for Organismal Studies (COS), University of Heidelberg, 69120 Heidelberg, Germany; HCI-IWR, Heidelberg University, 69120 Heidelberg, Germany; EMBL Heidelberg, 69120 Heidelberg, Germany

**Keywords:** Keywords Arabidopsis, lateral root, cell division, segmentation

## Abstract

Lateral root formation determines to a large extent the ability of plants to forage their environment and thus their growth. In *Arabidopsis thaliana* and other angiosperms, lateral root initiation requires radial cell expansion and several rounds of anticlinal cell divisions that give rise to a central core of small pericycle cells, which express different markers than the larger surrounding cells. These small central cells then switch their plane of divisions to periclinal, and give rise to seemingly morphologically similar daughter cells that have different identities and establish the different cell types of the new root. Although the execution of these two types of divisions is tightly regulated and essential for the correct development of the lateral root, we know little about their geometrical features. Here we analyse a four-dimensional reconstruction of the first stages of lateral root formation and analyze the geometric features of the anticlinal and periclinal divisions. We identify that the periclinal divisions of the small central cells are morphologically dissimilar and asymmetric. We show that mother cell volume is different when looking at anticlinal versus periclinal divisions and the repeated anticlinal divisions do not lead to reduction in cell volume although cells are shorter. Finally, we show that cells undergoing a periclinal division are characterized by a strong cell expansion. Our results indicate that cells integrate growth and division to precisely partition their volume upon division during the first two stages of lateral root formation.

## Introduction

Plants have devised efficient strategies to maximize their uptake of resources and adapt to changing conditions. Instrumental to this is the continuous post-embryonic formation of new organs. Branching of new roots from the embryo derived primary root is an important determinant of a plant root system architecture and of plant fitness (Motte et al. 2019). In *Arabidopsis thaliana*, formation of lateral roots starts with the selection of founder cells in the basal meristem from xylem pole pericycle cells that retain the ability to divide and later initiate lateral root formation in the differentiation zone (Moreno-Risueno et al. 2010; Smet et al. 2007). Divisions of these founder cells marks the initiation of the lateral root (Dubrovsky et al. 2001). Further divisions give rise to the lateral root primordium (LRP) that then grows into a fully functional lateral root as it emerges from the primary root (Banda et al. 2019). The ontogeny of the LRP has been described in detail (Malamy and Benfey 1997). Typically three to eight adjacent cell files, each containing mostly two founder cells, contribute to the formation of the LRP (Dubrovsky et al. 2000). Among these, one to two take a leading role dividing first and contributing most cells to the LRP. These cell files are named “master” cell files and recruit additional flanking cell files (Torres-Martínez et al. 2020; von Wangenheim et al. 2016). LRP formation is a self-organizing process driven by cell growth and division which, despite variation in the number of founder cells and absence of deterministic sequence of divisions, converges onto a robust stereotypical tissue organization that allows to identify typical developmental stages (von Wangenheim et al. 2016). In total, eight stages (I-VIII) exist (Malamy and Benfey 1997), characterized by typical number of cell layers. Stage I is one cell layer thick and results from several rounds of anticlinal division (where the axis is colinear to the shoot-root direction) of the founder cells. These divisions are asymmetric, and result in two small inner daughter cells flanked on each side by two large outer cells (Malamy and Benfey 1997). The small inner cells then typically divide periclinally (division axis normal to the primary root-to-shoot axis), producing a second layer of cells, to mark the start of stage II (Malamy and Benfey 1997). Anisotropic cell growth (Vilches Barro et al. 2019), licensed by the active yielding of the overlying tissue (Vermeer et al. 2014) and further anticlinal and periclinal divisions occur in stages II-VIII and establish the dome-shaped LRP (Lucas et al. 2013).

Asymmetric cell division (ACD) is a common feature of multicellular organisms, and in plants ACD produces distinct cell types and new organs, and maintains stem cell niches. Daughter cells of ACD have different fates and/or different volume. Because plant cells are encaged within the cell wall, the correct development of organs relies on strict coordination of asymmetric cell divisions in space and time (De Smet and Beeckman 2011). The asymmetric anticlinal division of the founder cells occurring at stage I is tightly controlled and entails a precise coordination of anisotropic cell growth and cytoskeletal reorganization (Vilches Barro et al. 2019), a situation reminiscent of the early embryogenesis (Kimata et al. 2016, 2019). This first asymmetric division of the founder cell is regulated by auxin signaling and ensures that it always produces two small inner cells and two larger outer cells (De Smet et al. 2008; Goh et al. 2012). The geometrical asymmetry of the division is obvious from the different length of each daughter cells, but as the founder cells expand anisotropically (Vilches Barro et al. 2019), it remains unknown how the volume is partitioned between the inner and outer daughter cells. Furthermore, several rounds of anticlinal divisions can occur, and it is unknown whether all these divisions have similar geometrical characteristics (De Smet et al. 2008). Interestingly, the number of these extra rounds of divisions can be modulated by the properties of the cell wall, hinting at a link between cell growth and the execution of the division (Ramakrishna et al. 2019). The transition from a stage I to a stage II LRP corresponds to the switch from anticlinal to periclinal divisions splitting the inner cells along their long axis, in violation of the shortest wall principle (Rasmussen and Bellinger 2018). Periclinal divisions produce daughter cells with different identities that contribute to different parts of the root primordium but it is unknown whether the first periclinal divisions are asymmetric and whether their execution entails specific geometrical features. Deviation from such geometric division has been reported to be driven by auxin in the early embryo (Yoshida et al. 2014). Interestingly, members of the PLETHORAs transcription factor family that mediates auxin effects, have been shown to specifically control the stage I to stage II transition as their mutation leads to the non-execution of the periclinal division and results in malformed LR primordia that fail to pattern (Du and Scheres 2017). These periclinal divisions produce daughter cells with different identities that contribute to different parts of the root primordium (Malamy and Benfey 1997), but it is unknow whether these first periclinal division are asymmetric and whether their execution entails specific geometrical features.

Here, we analyze in details all the divisions of the founder cells leading to a stage II LR, focusing on the cell volume. We used light sheet microscopy to obtain high resolution live imaging of wild type *Arabidopsis thaliana* LR primordia expressing cell contour and nucleic markers. Segmentation of the nuclei and of the cells in these images allow us to extract the geometric characteristics of each division and achieve a detailed analysis of the cell volume partition during divisions. We show that differences exist between the mother cells of anticlinal and periclinal divisions that the periclinal divisions marking the transition to stage II are themselves asymmetric. Consecutive rounds of anticlinal, although of seemingly different 2D geometry have similar volume and that periclinal divisions are preceded by intense cell growth. Our results suggest that cell growth and division are integrated by the LR cells which precisely partition volume upon division.

## Results

We combined live imaging with nuclei and cell segmentation to analyse how cells partition during divisions of LR founder cells (Figure 1A).

**Figure 1.**
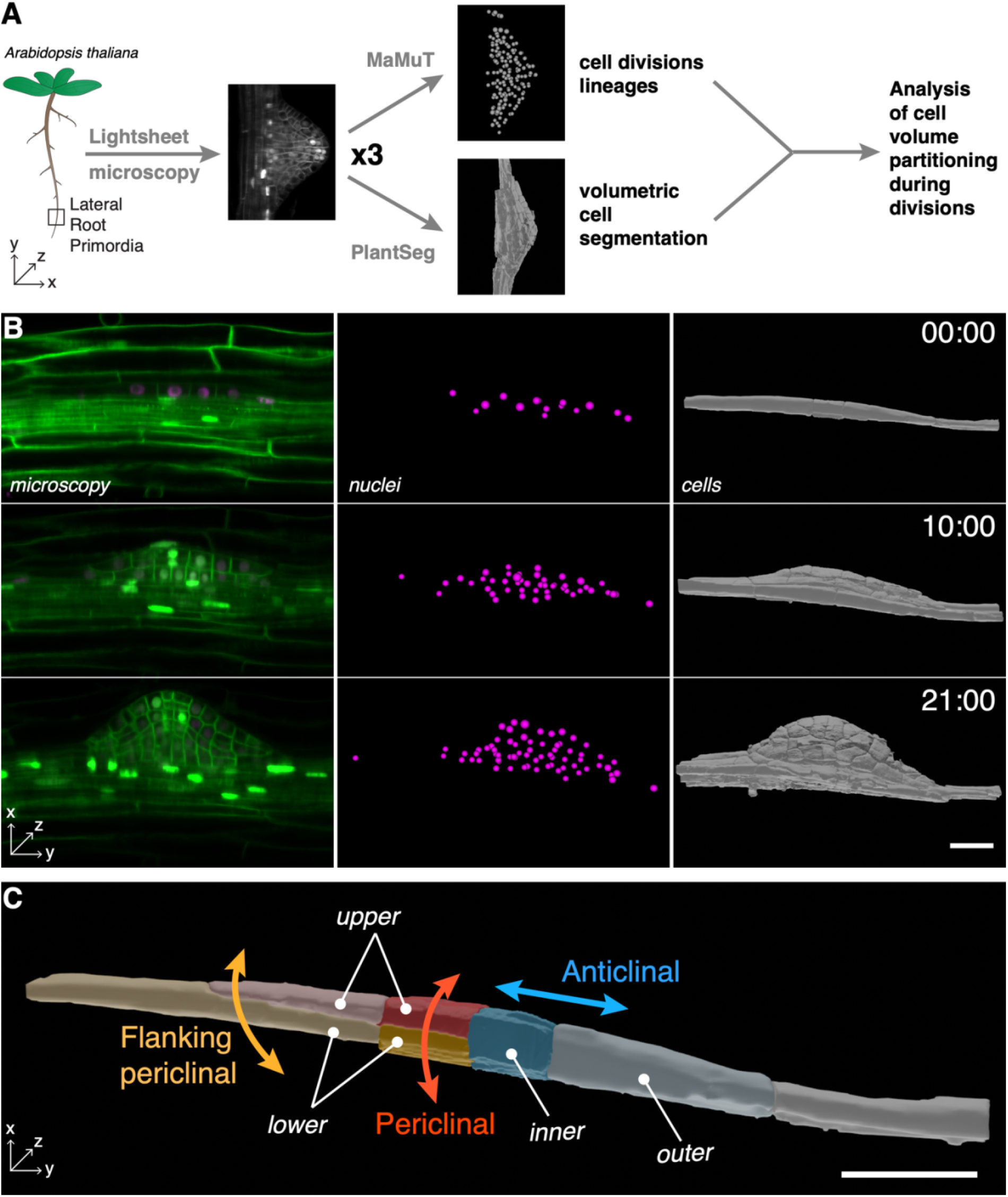
Analysis of cell volume partitioning during division. (A) Overview of the analysis. Three *Arabidopsis thaliana* lateral root primordia (LRP) are imaged by light sheet microscopy. Nuclei of the LRP are tracked by MaMuT and the divisions classified while the cells are segmented using PlantSeg. The merged data are then analyzed. (B) Snapshots of development of the LRP B. Left, a single median slice of the LRP imaged by light sheet microscopy visualized using the sC111 reporter (see methods). Center, visualisation of LRP nuclei position. Right, PlantSeg segmented cells. The time (hh:mm) of the recording is indicated. (C) Division types and relative daughter cells locations. Scale bars: 25μm.

### The datasets

We used light sheet fluorescence microscopy to capture the development of three lateral root primordia during 25 to 46h, imaging every 30mins. All three lateral root primordia, hereby called A, B & C (Table 1) presented a typical morphology and speed of development and were thus representative of lateral root formation. Development of the lateral root primordia A and B was recorded from stage I on. Primordium C and from early stage II for primordium C, where one cell had already divided periclinal, was recorded from early stage II on (Figure 1B, Figure S1, S2 and movies S1 to S3). As previously reported (Torres-Martínez et al. 2020; von Wangenheim et al. 2016), the spatial organization of the founder cells, and their contribution varied from primordium to primordium (Figure S1). We first identified cell files which contributed more to the development of the LRP, the so-called master cell files (Torres-Martínez et al. 2020; von Wangenheim et al. 2016). In primordium A, a single cell file contributed ~50% of cells to the primordium (total of ~ 90 cells, corresponding to stage V) and could be unambiguously labelled as the “master” cell file. In primordium B and C, no single cell file contributed to more than 42% of the cells, we therefore labelled as “master” two cells files with most important contributions (Figure S1). To confirm that these master cell files had indeed a pioneering role (von Wangenheim et al. 2016), we compared the timing of the transition from stage I to stage II. In all primordia, the first periclinal division marking the transition to stage II occurred 5h20min ± 1h (avg. ± sd, n = 3) earlier in the master cell files than in the flanking peripheral cell files (Table 1). This confirmed that the master cell files we identified lead the development of the primordia.

**Table 1.** Characteristics of the lateral root primordium analyzed.

| Primordium | Stage at t0 | Recording duration | Lineage tracked for | Segmented for | 1st Periclinal division in master cell file | 1st Periclinal division in peripheral cell file |
| --- | --- | --- | --- | --- | --- | --- |
| **A** | I | 39h | 20.5h | 20.5h | 7.5h | 11.5h |
| **B** | I | 25h | 20.5h | 20.5h | 2.5h | 9h |
| **C** | I/II | 46h | 31h | 31h | 1h | 4.5h |

### The analysis pipeline

First we established the complete cell lineage of each lateral root primordia with the Fiji/ImageJ plugin Multi-view Tracker (MaMuT), used to annotate cell behaviors in 4D (Wolff et al. 2018). Each lineage was followed until the first periclinal division or, if this never occurred, the final anticlinal division. Each cell division was classified as anticlinal (division axis colinear to the shoot-root axis and generating more cells in a given file) or periclinal (where the axis is normal to the surface main root and generates new layers) (Figure 1C). We further divided the periclinal division into “normal” and “flanking” as there were visible differences in shape and size (Figure 1C). Whereas the “normal” periclinal divisions were typically found in the center of the primordium and had a division plane perpendicular to anticlinal walls, the “flanking” ones were typically observed on the distal ends of the primordia and had a typical oblique orientation where the cell is split from the upper cell vertically on one end and horizontally on the other (Figure 1C). In addition to their type, we recorded additional metadata about these divisions (see methods). A total of 166 daughter cells for 83 divisions were analyzed and the results compiled in a single file.

Second, we used PlantSeg (Wolny et al. 2020) to segment all cells of the three lateral root primordia and overlying tissues. The segmented cells were visualized using the software Blender and segmentation errors (under-segmentation requiring to split cells and over-segmentation requiring to merge cells) were corrected (see methods). The volumes of the lateral primordia cells tracked with MaMuT were then retrieved from the curated segmentation and merged with the division data. We obtained volume information for 106 cells representing 53 divisions.

### A limited set of division sequences lead up to stage II

Stage I primordia result from anticlinal divisions that split a single or a pair of abutting LR founder cells (Torres-Martínez et al. 2020; von Wangenheim et al. 2016). The number of these divisions is variable from primordium to primordium (Ramakrishna et al. 2018; Torres-Martínez et al. 2020; von Wangenheim et al. 2016), but typically leads to a configuration with small central cells (inner) flanked by longer cells (outer). The central-most cells then reorient their division planes and undergo formative periclinal cell divisions to generate a new cell layer, leading to a stage II primordium, an essential transition for proper LR development (Du and Scheres 2017). We thus investigated whether there was any regularity in the sequence of anticlinal divisions leading to the switch to Stage II. For this, we analysed 28 complete division sequences until the first periclinal for founders located in central and peripheral cell files (Figure 2A). In all cases, the founder first divided anticlinal, producing a small inner cell and a larger outer cell. In most cases (100% n=13 in the master cell files and 87% n=15 in the periphery), the inner cells divided periclinal in the second round of division. In two cases, the inner cells underwent a second anticlinal asymmetric divisions producing a long outer cells and a smaller inner cell which invariably then divided periclinal. In comparison to this, most outer cells resulting from the first anticlinal division often divided anticlinal again in the second and third rounds of division or performed an oblique “flanking” periclinal division. Taken together, we could identify a re-iterated pattern, where after an asymmetric anticlinal division, the smaller cells switch to periclinal divisions while the larger cells either do a flanking periclinal division or divide again anticlinal and repeat the pattern (Figure 2B).

**Figure 2.**
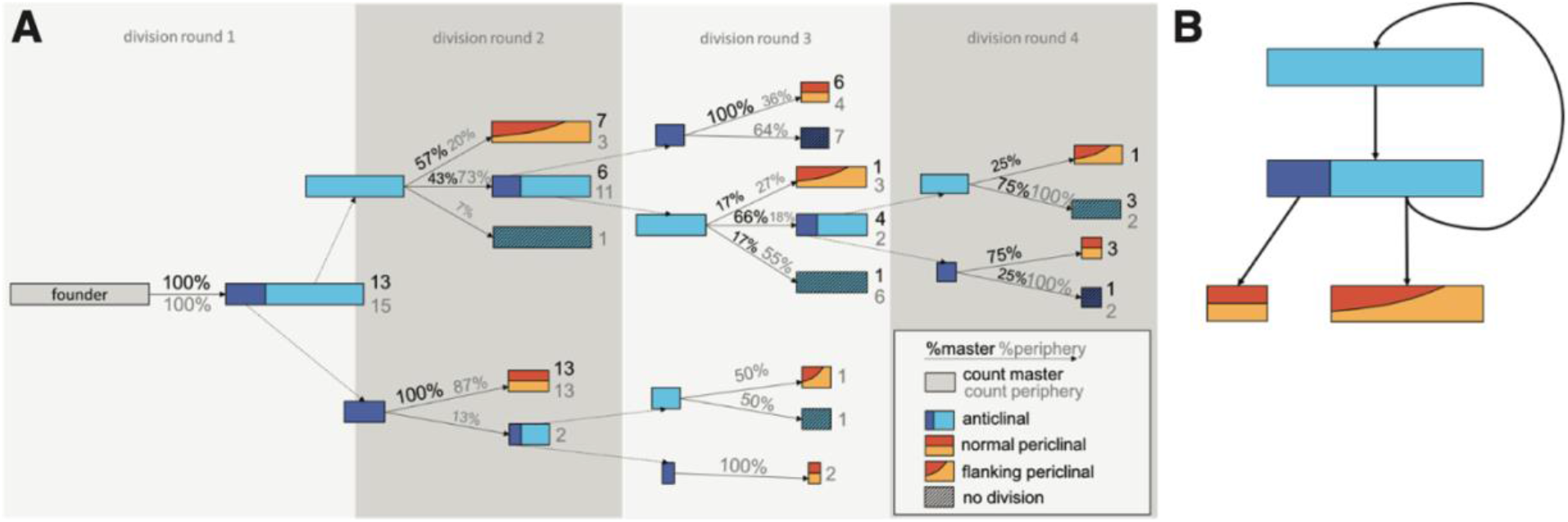
Division sequence leading to a stage II LRP. (A) Division sequences and frequencies of individual division events for master and peripheral cell files. (B) Re-iterated pattern during the first divisions of the LRP. Elongated cells undergo an asymmetric anticlinal division, producing a small cell that then undergoes a periclinal division and a large cell undergoing a flanking periclinal division. Alternatively, the large cell repeats the sequence.

### Anticlinal and periclinal divisions are asymmetric and differ in the volume of the mother cell

The existence of this iterated pattern of divisions suggests that the switch to a periclinal division type may require specific geometric features. We thus examined the geometry, volumes and volume repartition during divisions in the three lateral root primordia. First, we looked at the repartition of volumes between the daughter cells of anticlinal and periclinal divisions. Because of their specific features, flanking periclinal divisions were analyzed separately (see below). For anticlinal divisions, the inner daughter cell is smaller than the outer one, with median volumes of 1009 ± 300 μm^3^ (median ± MAD) and 2122 ± 1647 μm^3^, respectively. For periclinal divisions, the lower daughter cell (762 ± 213 μm^3^) is smaller than the upper one (1144 ± 392 μm^3^) (Figure 3A). This asymmetry in segregation of volume in the daughter cells is represented by a volume ratio (inner/outer for anticlinal divisions and upper/lower for periclinal divisions) lower than 1 for the two types of divisions: 0.59 ± 0.31 (median ±MAD) for anticlinal, 0.77 ± 0.40 for periclinal (Figure 3B). We then looked whether the decision for a cell to undergo an anticlinal or versus a periclinal division correlated with its volume. For this, we estimated the volume of the mother cells at the time of division as the sum of the volumes of the two daughter cells and plotted its distribution according to the type of division (Figure 3C). Cells undergoing anticlinal divisions are larger (3177 ± 2046 μm^3^) than the ones undergoing periclinal divisions (1795 ± 677 μm^3^). In addition to this difference in absolute volume, the coefficient of variation (cv = MAD/median – a measure of dispersion) reveals that mother cells of anticlinal divisions are more diverse in volume than those undergoing for periclinal divisions (cv_anticlinal_ = 0.64 vs. cv_periclinal_ = 0.37). Thus, the anticlinal and periclinal divisions leading to a stage II primordia are characterized by an asymmetric repartition of volume between daughter cells, and cells undergoing a periclinal division tend to be twice as small and more homogeneous in volume than the ones dividing anticlinal.

**Figure 3.**
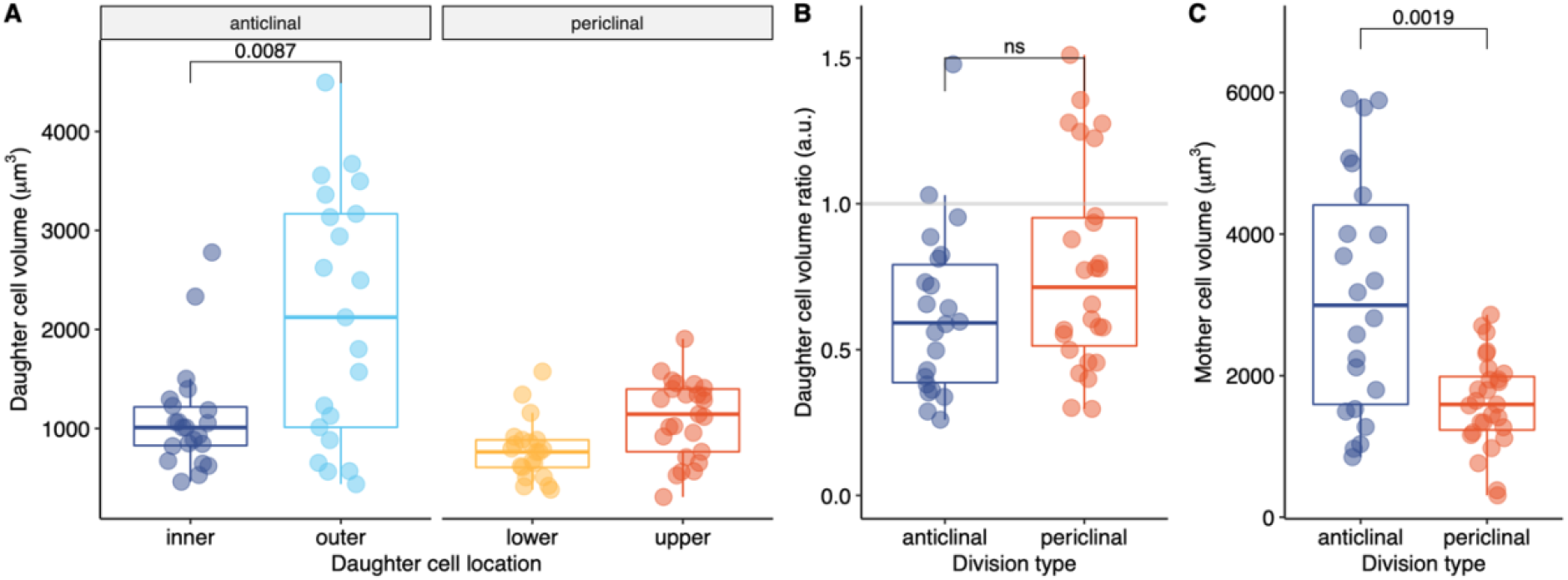
Volumes in mother and daughter cells of during anticlinal and periclinal divisions of all anticlinal and periclinal divisions observed in all primordia. (A) Distribution of Volumes of the inner and outer and lower and upper daughter cells for anticlinal and periclinal divisions (B) Distribution of the ratio of daughter cells volumes (inner/outer for anticlinal divisions and upper/lower for peri clinal divisions). (C) Distribution of the volumes of mother cells undergoing anticlinal or periclinal divisions. In all panels, flanking periclinal divisions are excluded. Comparison between pairs of samples was performed using the Wilcoxon rank-sum test and the p value indicated (not significant).

### Similar volume segregation in divisions occurring in the central and peripheral cell files

LRP development is characterized by the emergence of one or two master cell files that have pioneer roles and peripheral cell files that are subsequently recruited (Torres-Martínez et al. 2020; von Wangenheim et al. 2016). We asked whether differences existed in the distribution of volumes when divisions occurred in the master or the peripheral cell files. For this we examined the distribution of the volumes among the daughter cells of anticlinal and periclinal divisions taking place in the master or peripheral cell files (Figure 4). We did not observe any differences neither in the volume repartition (Figure 4A, B) nor in the ratio (Figure 4C). We thus conclude that the asymmetric nature of the divisions during the progression of the LRP from stage I to stage II is similar, no matter whether these occur in the master of in the later recruited peripheral cell files.

**Figure 4.**
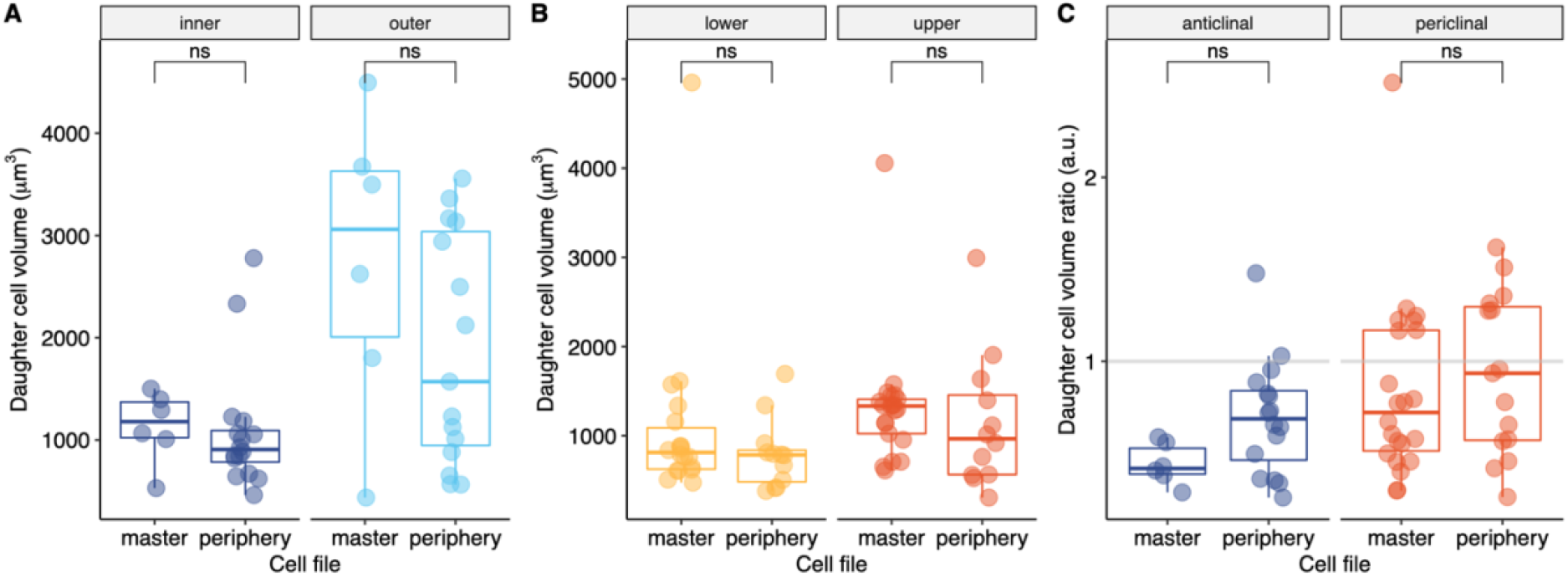
Daughter cells v olume s in master or peripheral cell files. Distribution of daughter cells volumes upon anticlinal (A) and periclinal (B) divisions as well as the ratio (C) for divisions occurring in the central or peripheral cells files. Comparison between pairs of samples was performed using the Wilcoxon rank-sum test and the p-value indicated (not significant).

**Figure 5.**
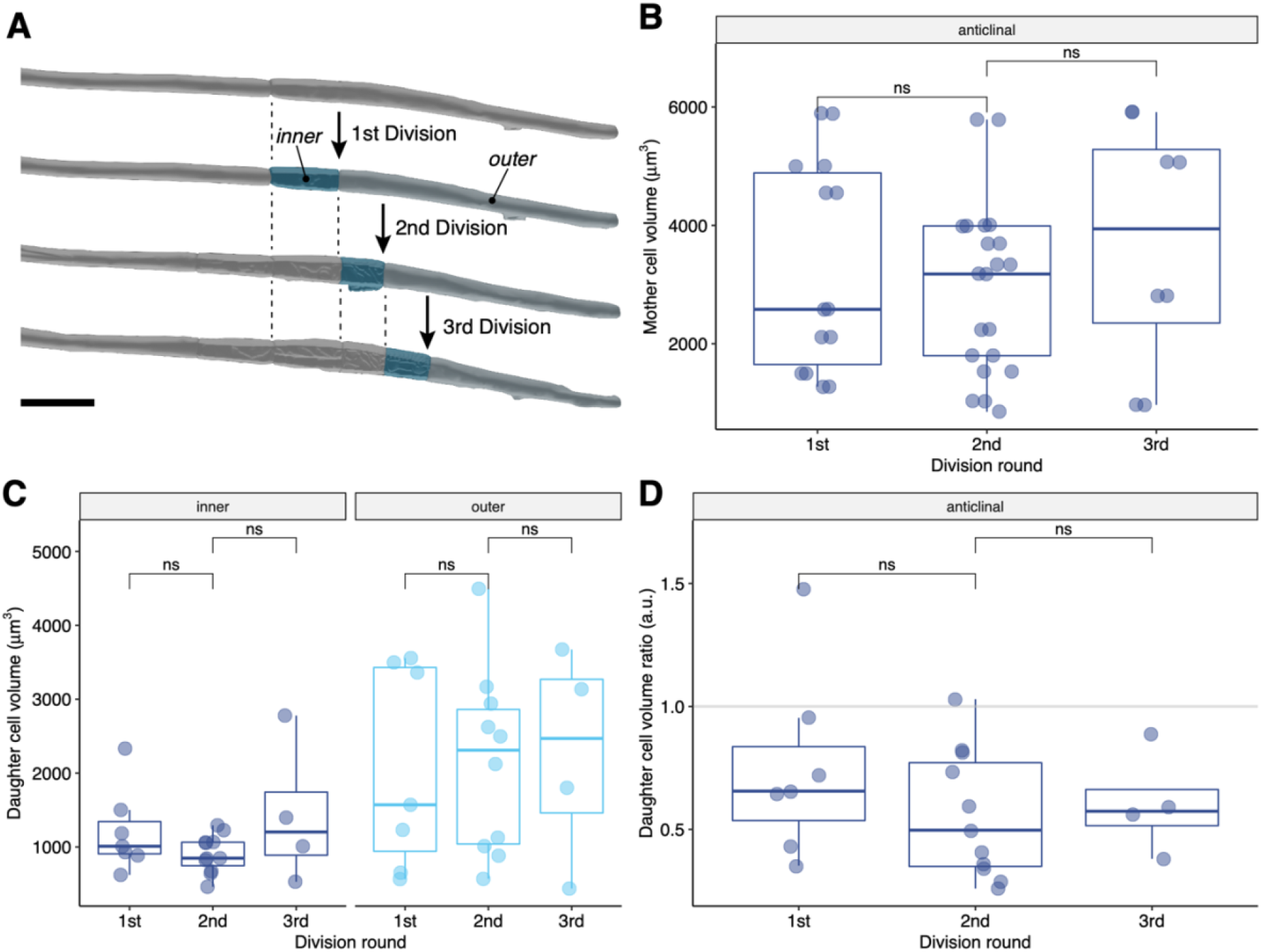
Volumes of daughter cells during consecutive rounds of anticlinal divisions. (A) Example of a founder cell undergoing three consecutive rounds of anticlinal divisions. Distribution of Volumes of the mother cells (B), of the inner and outer daughter cells (C) and their ratio (D) in three consecutive rounds of division. Comparison between pairs of samples was performed using the Wilcoxon rank-sum test and the p-value indicated (not significant).

### Consecutive division rounds are characterized by the same volume distribution between the daughter cells

LR founder cells can do several consecutive rounds of anticlinal divisions (Figure 2, 4A). This situation is interesting as it progressively shortens the outer cell and may change the volume repartition in the daughter cells. To investigate this, we first examined the mother cells that undergo consecutive anticlinal divisions (Figure 4A, B). We observed that the volume of these cells remains similar across all three division rounds (3177 ± 2046 μm^3^) although their length is reducing (from 75μm to 57μm, Figure S4), indicating that radial growth (Vilches Barro et al. 2019) is compensating the volume reduction induced by the consecutive divisions. The repartition of volume between the daughter cells of each consecutive round of anticlinal divisions is very similar both in absolute volume (Figure 4C) and in their ratio (Figure 4D). The ratios were constant at ~0.5, meaning the inner cell always received a third of the mother volume. Thus, anticlinal divisions are characterized by a characteristic absolute volume and distribution of among daughter cells. Together, this suggests that the combined effect of cell division and cell growth lead to similar volume partition in consecutive rounds of anticlinal divisions.

### Cells undergoing a periclinal division have a characteristic absolute volume and display more cell growth

The transition from stage I to stage II being a crucial for LR development and specifically controlled (Du and Scheres 2017), it is still unknown whether the stereotypical change in division plane is an automatic event due to the size or geometry of the small inner cells or whether it is controlled by molecular mechanisms that do not rely on size or shape. We previously saw that cells dividing periclinal have an smaller absolute volume than the ones dividing anticlinal. If a certain volume is necessary for the shift in division planes, then this should be maintained in all division rounds. We thus investigated the distribution of volume of inner cells resulting from consecutive rounds of anticlinal divisions and now undergoing periclinal divisions. (Figure 6A). There are no differences in volume (Figure 6B) across the subsequent rounds of division nor in the ratio of daughter cells volumes after a periclinal division (Figure 6C). The mother cells had a volume of 1618 ± 562 μm^3^ and the ratio 0.71 ± 0.34, meaning the upper cells receive ~64% and lower cells receive 36% of the mother volume. Thus, the shift to a periclinal division correlates with a specific maximal volume . Cell growth in the LR being anisotropic and more pronounced in the central and apical region (von Wangenheim et al. 2016) we investigated whether inner cells transitioning to a periclinal divisions had an increased cell expansion compared to cells dividing anticlinal. For this we first computed the difference between the volume of a mother cell (sum of the volume of the two daughter cells) and the volume of that same cell right after its last division. This difference, which measures how much the cell expanded between two divisions, is divided by the volume of the cell right after its last division to obtain a ratio of expansion:

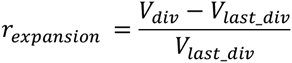

**Figure 6.**
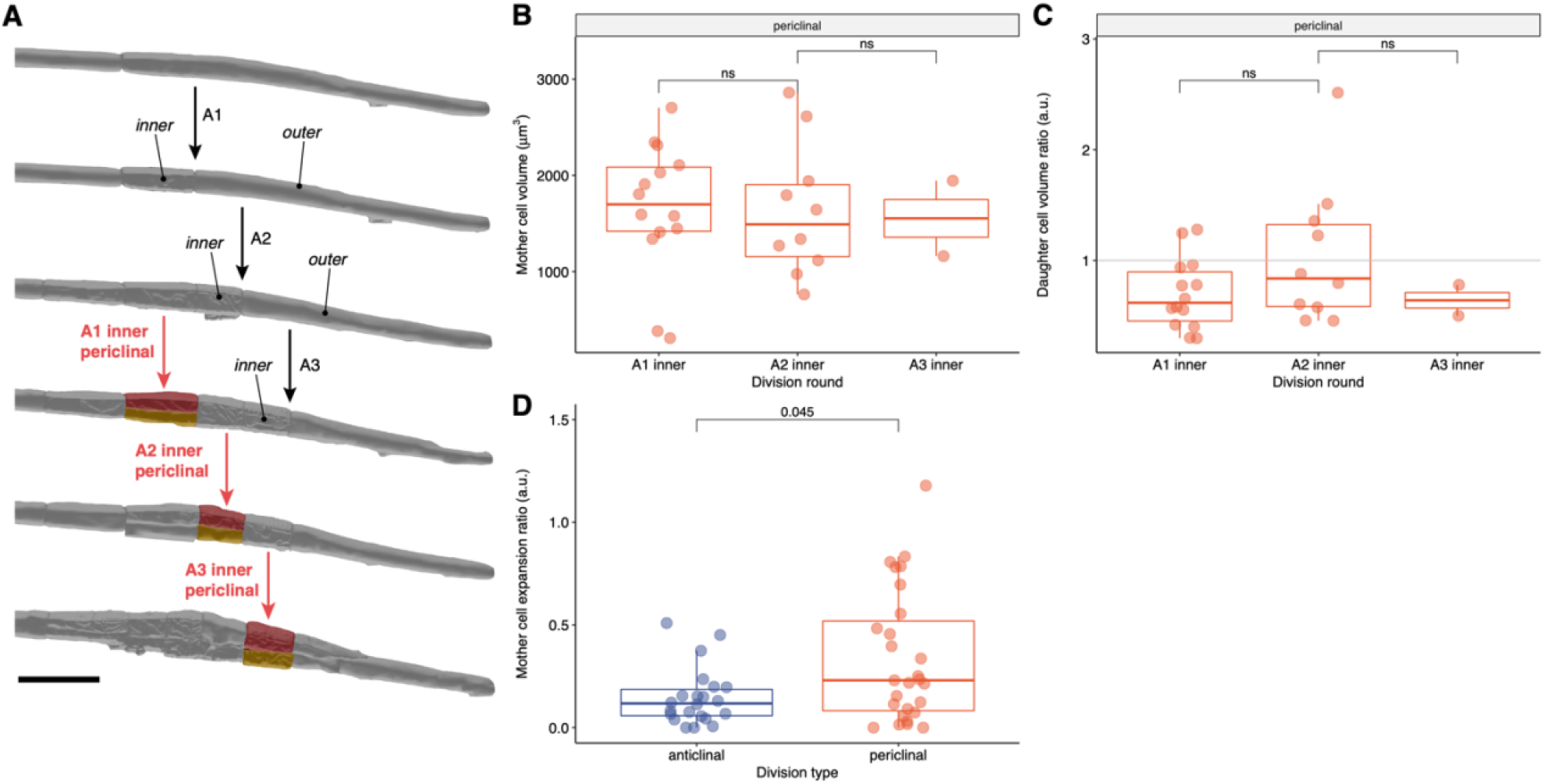
Volumes repartition and cell growth preceding periclinal divisions. (A) Periclinal divisions of inner cells resulting from three rounds of anticlinal divisions. Distribution of Volumes of the mother cells (B) and of the inner and outer daughter cells ratios (C) in three consecutive rounds of divisions. (D) Distribution of the ratio of mother cell expansion before anticlinal or periclinal division. Comparison between pairs of samples was performed using the Wilcoxon rank-sum test and the p-value indicated (not significant).

We plotted the distribution of this ratio for cells undergoing anticlinal of periclinal divisions. Mother cells of periclinal divisions (flanking periclinal divisions excluded) had more pronounced increase in growth since the last division 0.23 ± 0.29 (median ± MAD) compared to anticlinal ones (0.11 ± 0.1, p = 0.045 Wilcoxon rank test). Taken together, the transition to a periclinal division occurs in rapidly expanding inner cells with a volume of ~1600 μm^3^, indicating that specific geometric constraints exist on for the execution of this switch.

### Anticlinal divisions cannot be distinguished from flanking periclinal divisions based on mother cell sizes

The outer daughter cell of an anticlinal division can either enter another round of anticlinal division or switch to an flanking periclinal division that displays an oblique division plane (Figure 7A). We investigated whether this transition was associated with specific features in term of absolute mother cell volume or volume repartition. For this, we examined the distribution of outer cells that undergo either another round of anticlinal division (“anticlinal”) or switch to a periclinal division (“flanking periclinal”). We could not identify any differences in absolute volume or ratio of daughter cells volume between the two types of divisions (Figure 7B, C). Although not significant, we noticed that cells undergoing a flanking periclinal division were smaller (1796 ± 1236 μm^3^ vs. 3075 ± 2263 μm^3^ for anticlinal) and had a less variable volume distribution (cv_med_ = 0.68 vs. 0.73 for anticlinal). We then looked at the rate of growth of the outer cells between two divisions. The distribution of the expansion ratio rexpansion reveals that mother cells of flanking periclinal divisions had a more pronounced increase in growth since the last division 0.61 ± 0.36 (median ± MAD) compared to anticlinal ones (0.12 ± 0.1, p = 0.00026 Wilcoxon rank test). Together, outer cells switching to a flanking periclinal division have an absolute volume similar to the ones undergoing another round of anticlinal divisions but are characterized by a higher rate of interphasic expansion, similarly to the inner cells switching to periclinal division.

**Figure 7.**
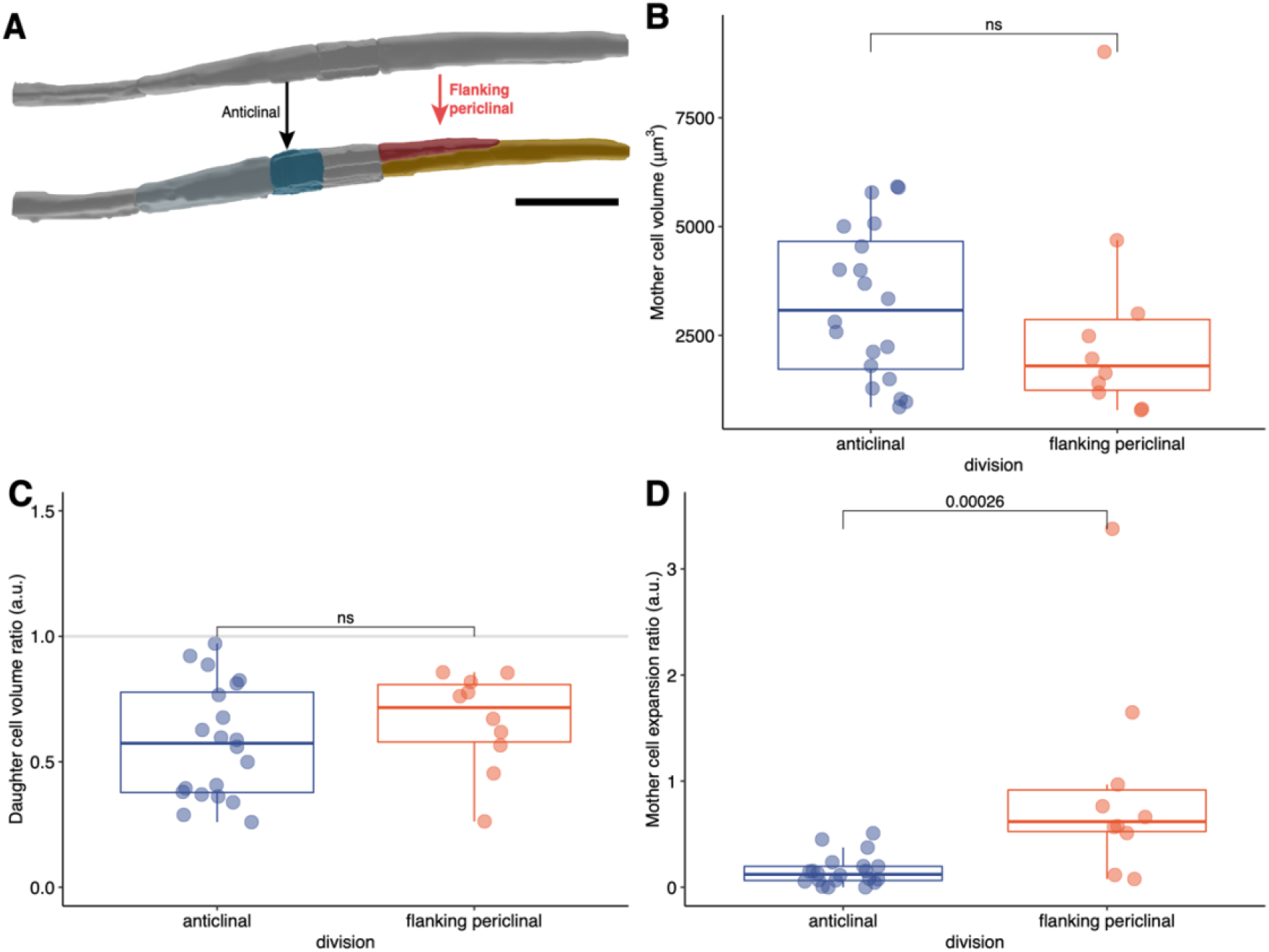
Volumes in daughter cells of flanking periclinal divisions. (A) Example of long daughter cells resulting from the switch of an anticlinal to a flanking periclinal division (right red arrow) or undergoing an extra anticlinal division (left black arrow). Distribution of Volumes of the mother cells (B), of the daughter cells ratios (C) or of the mother cell expansion ratio (D) in both cases. Comparison between pairs of samples was performed using the Wilcoxon rank sum test and the p value indicated (not significant).

## Discussion

We processed high resolution time resolved volumetric images of three Arabidopsis lateral root to classify all divisions occurring up to stage II and combine these information with the cell geometry derived from the volumetric segmentation of each cells. These three digital lateral roots allow us to get an unprecedented 3D look at the cellular architecture of the developing lateral root primordia.

Such digital reconstructions are useful tools to quantify cell and tissue behaviors during morphogenesis in plants. They allow precise quantification of the geometrical attributes of cells in complex tissues (Fernandez et al. 2010; Kierzkowski et al. 2012; Ripoll et al. 2019; Sapala et al. 2018; Vijayan et al. 2020) and when they are time resolved allow the inference of growth direction and intensities (Hervieux et al. 2016; Kierzkowski et al. 2012, 2019). Lateral root morphogenesis has been, with few exceptions (Lucas et al. 2013; Torres-Martínez et al. 2020; von Wangenheim et al. 2016), essentially studied using (optical) 2D sections. Although extremely valuable the lack of volumetric information can lead to wrong conclusions, especially when cells have non simple 3D geometries. Here, our analysis revealed that the periclinal divisions of the central cells which were thought to be geometrically symmetrical, give rise to daughter cells of markedly different volumes. This geometrical asymmetry was only revealed by a quantification of the 3D volume of the daughter cells. In 2D these divisions appear to split equally the cells in the middle but because of their trapezoidal shape, the upper cells are larger than the lower one. This is reminiscent of the case of the formative divisions occurring in the early Arabidopsis embryo that appear symmetrical in 2D but reveal asymmetrical in 3D (Yoshida et al. 2014). This geometrical asymmetry is in agreement with the formative nature of these divisions that initiate the formation of the different tissues of the lateral root (Goh et al. 2016; Malamy and Benfey 1997) and reinforce the view that in plants, asymmetries in volume partition among daughter cells and differential fate seem in many instance intertwined (De Smet and Beeckman 2011).

The several rounds of anticlinal divisions that lead to the formation of a central region with small cells flanked by larger cells have a characteristic partition of volume among daughter cells (~1/3 for inner daughter and ~2/3 for the outer one) and result from a reiterated motif. The founder cell divide asymmetrically to give rise to an outer larger daughter which in turn divides several times to give rise to small inner cells. Although the outer cells become progressively shorter, the volume of the cells that accomplish additional anticlinal division remains stable and the volume distribution among daughter remains also constant. This indicates that the radial expansion of the lateral root cells during stage I (Vermeer et al. 2014; Vilches Barro et al. 2019) counterbalances the shortening of the cells. It is interesting to note that mutation of *EXPANSIN-A1,* a gene encoding a cell wall remodeling protein, perturbs the capacity of the pericycle to radially expand and leads to extra rounds of anticlinal divisions at stage I and partition of volume among daughter cells (Ramakrishna et al. 2018). Similarly, perturbing the anisotropic expansion of the LR founder cells by interfering with the cytoskeleton also leads to mispositioning of the plane of division (Vilches Barro et al. 2019). Together, these suggest that cell growth (through remodeling of cell wall properties) and cell division might be homeostatically maintained and ensure a defined distribution of cell volume among daughter cells.

Periclinal division of the central cells marks the transition from stage I to stage II, a symmetry breaking event that establish new radial and proximodistal for the new LRP. This switch division orientation is controlled by specific regulators such as PLT3, PLT5 and PLT7 (Du and Scheres 2017) and initiate the expression of genes that mark the segregation of proximal–distal domains with expression patterns that differ between the inner and outer layers. We observe that this transition is characterized by two features. First, periclinal divisions occurs in cells with a volume of ~1800μm^3^ and this population is relatively homogenous while anticlinal divisions are typically observed in larger cells with a wider distribution of volume. This may suggest that only cells of a maximal volume and/or geometry may be competent for the execution of a periclinal division. What element could be responsible for this specific competency is speculative. Their typical geometry may lead to preferential accumulation of auxin as it is the case in the embryo (Wabnik et al. 2013), as supported by the higher auxin signaling in these small inner than the flanking one (Benková et al. 2003). High auxin and PLTs might thus contribute to the switch in division orientation. The second characteristic is that these central cells are characterized by an important rate of cell growth before the periclinal division. The central domain of the primordium is indeed the area of important anisotropic growth (Vilches Barro et al. 2019; von Wangenheim et al. 2016). Growth anisotropy and division orientation are linked (Sablowski 2016), the axis of the periclinal division may follow the principal direction of growth and/or align with the mechanical stress resulting from the anisotropy of growth (Louveaux et al. 2016). The role of cell expansion and possibly of mechanical cues in the control of division orientation is also visible for the flanking periclinal divisions. These divisions are only observed in the long cells on the flanks of the primordium and have atypical oblique division planes. Cell volume alone does not seem to determine this divisions as cells of similar volume undergo anticlinal divisions. Yet like in the case of the central cells, important cell expansion precedes a flanking periclinal division. It would be interesting to monitor the orientation of microtubules in these cells to allow the inference of direction of stress and growth direction (Uyttewaal et al. 2012).

## Materials and Methods

### Plant material and growth

Three *Arabidopsis thaliana* seedlings expressing the *UB10pro::PIP1,4-3xGFP / GATA23pro::H2B:3xmCherry / DR5v2pro::3xYFPnls / RPS5Apro::dtTomato:NLS* (sC111) reporter (Vilches Barro et al. 2019; Wolny et al. 2020) were used for imaging of LRP formation. Seedlings were sterilised and deposited on top of capillaries (100 μL, micropipettes Blaubrand Cat.-N° 708744) filled with ½MS medium containing 1% Phytagel (SIGMA-ALDRICH, Cat.-N° 71010-52-1), stratified at 4°C for 48h and grown 4-5 days at 22° C with light intensity of 130-150 μE/m^2^/sec and photoperiod 16 h /8 h day-night.

### Light sheet fluorescence microscopy

Imaging was done on a Luxendo Bruker MuViSPIM. The phytagel rod containing the seedlings were carefully pushed out from the capillary until only the root tip remained inside. The capillary was positioned in the microscope sample holder, the imaging chamber The cotyledons could therefore float on the liquid ½MS inside the chamber and be exposed to air. Imaging conditions and post-acquisition processing steps have been reported in (Wolny et al. 2020).

### Lineage tracking

Cell were annotated in 4D using the Fiji/ImageJ plugin Multi-view Tracker (MaMuT) (Wolff et al. 2018). For each time point, the image stacks corresponding to the nuclei signal were exported as one XML/HDF5 file pair using the Fiji/ImageJ plugin BigDataProcessor (Tischer et al. 2020) and annotated with MaMuT, with the integrated BigDataViewer (Pietzsch et al. 2015). For each nucleus its position (x,y,z,t) was recorded and it was linked to its future self at each consecutive time point, and, when cells divided, each daughter was linked to its mother. All divisions preceding and including the first periclinal division were manually annotated to include the type of division, the respective daughter cell locations, the time point, mother identity and the following division. All data were aggregated in a .csv file.

### Segmentation

The image stacks corresponding to the cell contours were cropped to only include the lateral root primordia and segmented using PlantSeg (Wolny et al. 2020) using the following parameters: CNN model: lightsheet_unet_bce_dice_ds1× with 100×100×100 patch segmentation algorithm: GASP with ß parameter of 0.65 and minimal size of 50000. For each time point each resulting segmented cell is identified by an unique label. Correspondence between these label and the nuclei tracked with MaMuT was established with a custom python script matching the position of the nuclei to the a segmentation label. Each segmented cell was converted to a mesh using a marching cube algorithm and exported as a PLY file with another custom script. Finally all meshes were imported into Blender (www.blender.org) for visualization, curation (splitting / merging of cells), annotation and volume calculation using a custom add-on called MorphoBlend and volumes exported to a .csv file. The two custom scripts are part of the PlantSeg-Tool toolbox which will be described, along MorphoBlend, in another manuscript. Meanwhile these tools can be obtained upon request.

### Data analysis

The lineage tracking and segmentation data were aggregated in a single data frame using R version 4.0.2 (2020-06-22) (R Core Team 2018) using the tidyverse (Wickham et al. 2019). All analysis and plotting were done in R using the tidyverse, ggpubr (Kassambara 2020) and ggsignif (Ahlmann-Eltze 2019) packages. Due to the lack of normal distributions, as determined by Shapiro-Wilk tests for normal distribution, Kruskal-Wallis tests were used to compare multiple independent groups and Wilcoxon rank-sum tests were used to compare two independent groups. The median was used, due to the non-normal distribution and the large number of outliers, to descriptively compare groups. Thus, the median absolute deviation (MAD) was used as the measure of variance. The source data and a R notebook describing all steps of the analysis is provided as supplemental material.

## ACKNOWLEDGEMENTS

We thank Marisa Metzger for her help in the preparatory phase of this project.

## AUTHORS CONTRIBUTION

ML and AM designed the study. ML generated the light sheet datasets. LMS and ML generated the lineages data and classification of divisions. LC, AW and AM wrote the scripts for processing of PlantSeg results and the visualization in Blender. AVB, SB and AM curated the segmentation results. LMS, ML and AM performed the analysis of the data. AK, FAH and AM interpreted the data. AM wrote the manuscript with input from all other authors.

## SUPPLEMENTAL MATERIAL

- Supplemental movies S1 – S3 Development, nuclei tracking and volumetric cell segmentation of the three lateral root primordia analysed.
- Figure S1. Frontal view of the organization of four LRP at timepoint 0 of each movie
- Figure S2. Side view of the master cell files of the four LRP at timepoint 0 of each movie.
- Figure S3. Volumes in daughter cells of anticlinal periclinal divisions in each LRP.
- Figure S4. Length of inner and outer daughter cells in consecutive rounds of anticlinal divisions.
- Supplemental file 1. CSV file of the106 daughter cells analysed.
- Supplemental file 2. CSV file containing the metadata about the 106 daughter cells analysed.
- Supplemental data 3. R notebook detailing the analyses performed.
- Supplemental data 4. PDF version of the R notebook.

