## Supplemental material for "Integration of Cell Growth and Asymmetric Division During Lateral Root Initiation In *Arabidopsis thaliana*"

**Supplemental figures and legends**

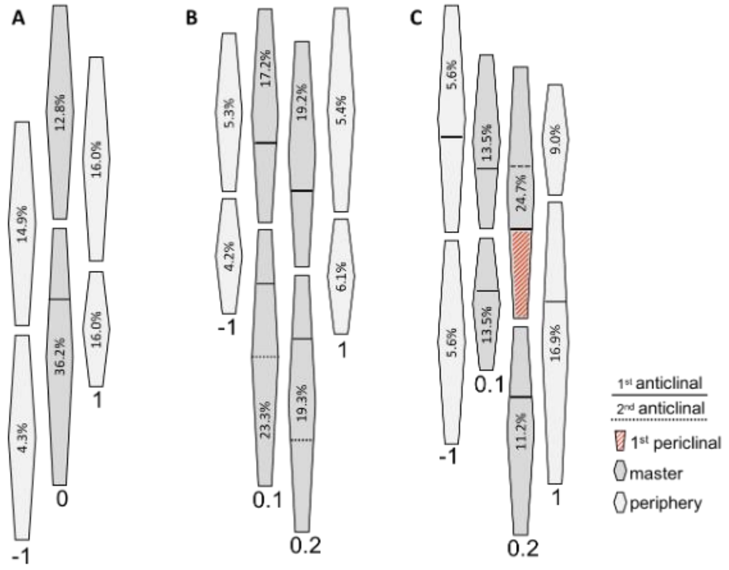


**Figure S1. Frontal view of the organisation of four LRPs at timepoint 0 of each movie**. Percent indicate the the contribution of each cell file in the 90-cell primordium. Single master cell files are given the index 0 while two neighboring master cell files are indexed 0.1 and 0.2. Peripheral cell files to the right (frontal view) are indexed 1 to n, and to the left -1 to -n.

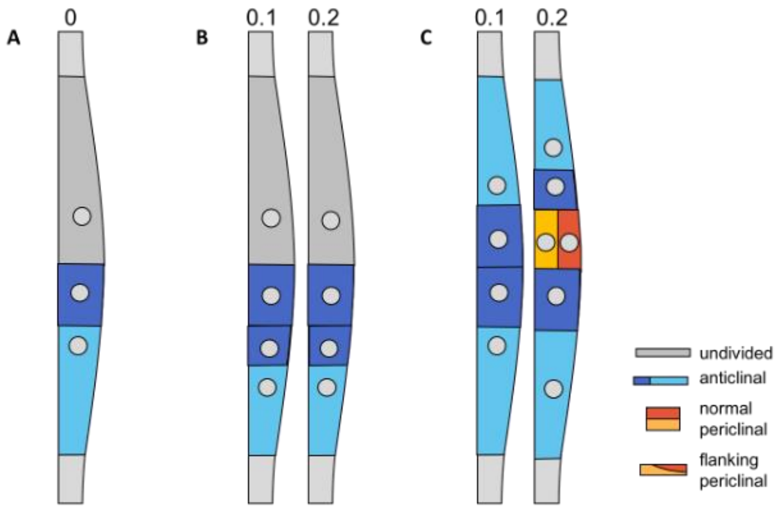


**Figure S2. Side view of the master cell files of the four LRPs at timepoint 0 of each movie.**

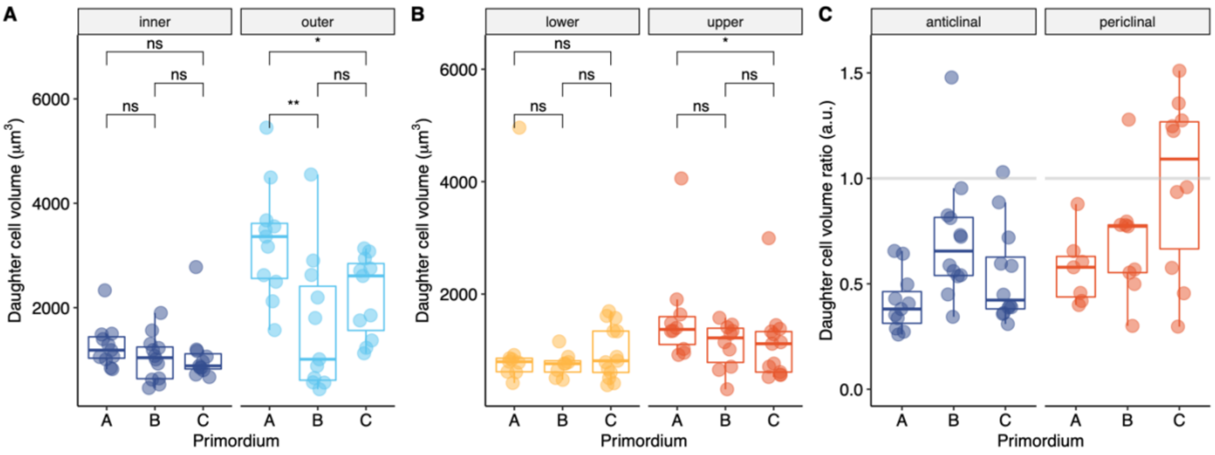


**Figure S3. Volumes in daughter cells of anticlinal periclinal divisions in each LRP.** Distribution of daughter cells volumes of anticlinal (A), periclinal (B) divisions or their ratios (C) for each LRP analyzed. Comparison between pairs of samples was performed using the Wilcoxon rank-sum test and the p-value indicated (ns. not significant).

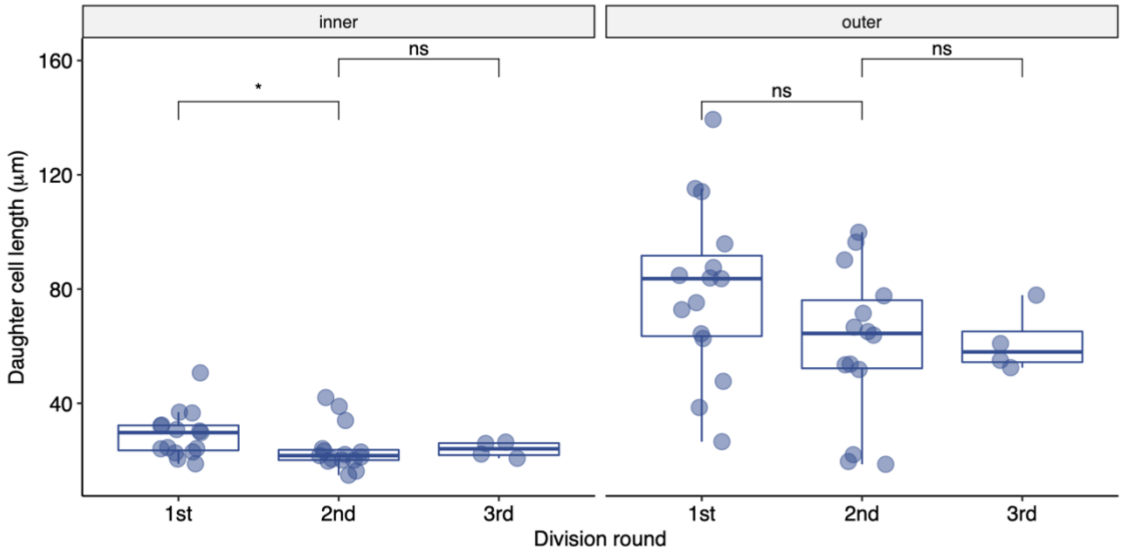


**Figure S4. Length of inner and outer daughter cells in consecutive rounds of anticlinal divisions.** Distribution of daughter cells length (along the root / shoot axis) of inner and outer cells. Comparison between pairs of samples was performed using the Wilcoxon rank-sum test and the p-value indicated (ns. not significant).
